# The Arabidopsis gene *RGO* mediates cytokinin responses and increases seed yield

**DOI:** 10.1101/2020.02.06.937342

**Authors:** Jhadeswar Murmu, Ghislaine Allard, Denise Chabot, Eiji Nambara, Raju Datla, Shelley Hepworth, Rajagopal Subramaniam, Jas Singh

## Abstract

A novel gene, *At1g77960*, from *Arabidopsis thaliana* was characterized. *At1g77960* transcripts accumulate to very high levels in plants ectopically overexpressing the *Golden2-like1* (*GLK1*) transcription factor and is designated as a *<u>R</u>esponse to <u>G</u>LK1 <u>O</u>verexpression* (*RGO*) gene. *RGO* encodes a protein with domains of tandem QH and QN repeats. Transcripts and promoter GUS reporter analyses indicated that *RGO* is expressed in roots, leaves, stems, floral and siliques tissues but not in seeds. Expression of the RGO:YFP fusion protein demonstrated that *RGO* is localized to the endoplasmic reticulum. MicroRNA mediated silencing of *RGO* resulted in severe reductions in vegetative and root growth, delayed flowering and reduced seed yield and viability, suggesting that *RGO* is essential for plant development. Conversely, ectopic overexpression of *RGO* resulted in enhanced vegetative growth including increased axillary bud formation and a 20% higher seed yield. Stable overexpression of *RGO* in *Brassica napus* also produced a similar increase in seed yield. Cytokinin (CK) response assays including root growth, green calli formation from excised hypocotyls and chlorophyll retention during dark-induced senescence suggest that one role of *RGO* is to mediate CK responses in plant development. These results suggest that *RGO* could be a target gene for increasing crop seed yields.

**One-sentence summary:** *RGO*, a novel gene from Arabidopsis, is essential for plant development, mediates CK signaling and increases seed yield in Arabidopsis and rapeseed when overexpressed.

## Introduction

Significant progress have been made in our understanding of the role of cytokinins (CK) in plant growth, and development. Through interactions with other hormonal pathways and transcriptional factors, CKs plays an important role in a variety of plant processes including greening (Cortleven and Schmülling, 2015), etioplast-to-chloroplast transition (Cortleven et al., 2016), root greening (Kobayashi et al., 2012; 2017), shoot and shoot development (Howell et al., 2003; Kieber and Schaller, 2014; Raines et al., 2016) and delay in leaf senescence (Gan and Amasino, 1995; Kim et al., 2006; Kieber and Schaller, 2014; Raines et al., 2016; Talla et al., 2016). The current knowledge of CK signalling involves a multistep signal transduction system that begins with the perception of CK by the Arabidopsis Histidine Kinases (*AHK2*, *AHK3*, *CRE1*/*AHK4*) in the endoplasmic reticulum (ER) phosphate transfer via the Arabidopsis Histidine Phosphotransfer proteins (AHPs). The information is transmitted to the nucleus where the transcription factors such as the type-B Arabidopsis Response Regulators (ARRs) affect gene expression (Kieber and Schaller, 2014; Romanov et al., 2018).

The roles of CK in chloroplast development are mediated by core components of the CK signalling pathway which includes the CK receptors *AHK2* and *AHK3* and the response regulators *ARR1*, *ARR10* and *ARR12* (Cortleven and Schmülling, 2015). Chloroplast development and maintenance in leaves also requires functional *Golden2-like* (*GLKs*) transcription factors (Fitter et al., 2002; Yasumura et al., 2005; Waters et al., 2008; Waters et al., 2009) which are members of the Myb superfamily of transcription factors containing a DNA binding “GARP” domain named after the maize *<u>G</u>olden2*, the type-B *<u>A</u>rabidopsis Response Regulators* (*ARR*), and the Chlamydomonas *<u>P</u>hosphate starvation response* (*PSR1*) transcription factors (Reichmann et al., 2000). In Arabidopsis leaves, *GLK1* and *GLK2* operate in redundant fashion as only the *glk1 glk2* double mutant showed impaired chloroplast development to display a distinct pale green phenotype (Fitter et al., 2002). The role of *GLK*s in chloroplast maintenance is also linked to leaf senescence, underscored by the observation that elevated expression of the transcription factor *ORE1* triggered early senescence in Arabidopsis through down regulation of *GLKs* and it has been observed that overexpression of *GLK1* delays leaf senescence (Rauf et al., 2013; Garapati et al., 2015). The role of *GLKs* in leaf senescence is also reflected in the observation that overexpression of *GLK1* resulted in the reduction of infection of the necrotrophic pathogen *Botrytis cinerea* (Murmu et al., 2014) as it is known that delay of senescence is a key factor in resistance to this pathogen (Swartzberg et al., 2008; Lai et al., 2011; Wang et al., 2013; Haffner et al., 2015).

A direct link between *GLKs* and CK signalling however, is lacking. There is evidence to suggest that type B *ARRs* coordinate the expression of *HY5* (elongated hypocotyl 5), which is required by *GLK2* for maximal (although not essential) greening in roots (Kobayashi et al., 2012). Overexpression of *GLK2*, *GNC (GATA*, *NITRATE-INDUCIBLE*, C*ARBON-METABOLISM-INVOLVED*) and *CGA1* (*CYTOKININ-RESPONSIVE GATA 1/GNC-LIKE)* result in root greening, however, neither *GLK1* nor *GLK2* are required for the CK-dependent root greening phenotype (Kobayashi et al., 2012). In tomato, overexpression of *GLK2* in the fruit leads to enhanced CK responsiveness and delayed ripening, thus differentiating function between the two *GLKs* in development programs such as fruit ripening (Lupi et al., 2019). However, the role of *GLKs* in CK response is lacking in leaves. There is no evidence to suggest that *GLKs* can participate in the phosphor-relay step and in this regard, are similar to the CK response regulator *ARR21C*, a truncated form of *ARR21* where the phosphor-relay N-terminus domain is removed (Kiba et al., 2005). Ectopic overexpression of *ARR21C* resulted in altered CK signalling, affecting plant development. *ARR21* has been shown to function in CK signalling as it can complement *arr1* and *arr12* mutants, although its precise role in CK signalling remains to be established (Hill et al., 2013). With the loss of the phospho-regulatory domain, ectopic overexpression of *ARR21C* in Arabidopsis produced a range of highly abnormal phenotypes with hypersensitivity to exogenous applications of very low levels of CK (Tajima et al., 2004; Kiba et al., 2005). To explore if an association exists between *GLK1* overexpression and CK signalling, we focussed on a previously uncharacterized Arabidopsis gene *At1g77960*, which was highly upregulated by ectopic overexpression of *GLK1* (Savitch et al., 2007). An examination of the microarray data from CK hypersensitive plants overexpressing *ARR21C* (Goda et al., 2008) also indicated that this gene was also substantially overexpressed.

Few other studies have implicated *At1g77960* in different plant processes. The *At1g77960* gene transcripts was upregulated in the *esr1-1* (enhanced stress response 1) mutant with increased resistance to the fungal pathogen *Fusarium oxysporum* (Thatcher et al., 2015) and was downregulated after infection with the bacterial pathogen *Pseudomonas syringae* pv tomato Dc3000 (Lewis et al., 2015). Despite the correlation between levels of gene expression in various plant processes, detailed characterization of *At1g77960* is lacking. Therefore, in this study we designated *At1g77960* gene as *<u>R</u>esponse to <u>G</u>LK1 <u>O</u>verexpression (RGO)*, and investigated its potential role in modulation of CK response and its significance in plant development, performance and seed yields.

## Results

### Arabidopsis *RGO* is a novel gene discovered from ectopic overexpression of *GLK1*

Transcripts of *AtRGO* were observed to be highly accumulated in Arabidopsis plants ectopically overexpressing *GLK1* (Savitch et al., 2007). A detailed RT-PCR and qPCR analyses demonstrated that *RGO* transcripts are highly upregulated in plants constitutively overexpressing *GLK1* (Fig. 1, B and C). Transcript levels in the *glk1* mutant remained unchanged when compared to the WT (Fig. 1, B and C) suggesting that *RGO* is not directly regulated by *GLK1*. An examination of the microarray data from CK hypersensitive plants overexpressing *ARR21C* also upregulated *RGO* transcripts by nine fold (Supplementary Table. S1). To further substantiate the observation of *GLK1* in the activation of *RGO* overexpression, we cloned and overexpressed *TaGLK1* from bread wheat (NCBI Accession EF105406) in Arabidopsis. Arabidopsis plants overexpressing *TaGLK1* also showed a high upregulation of *RGO* (Supplemental Fig. S1). *RGO* is a single copy gene encoding a 48 kilo Dalton (kDa) glutamine (Q) rich (14%) protein of 420 amino acids. The N-terminus of the deduced protein contains a hydrophobic region (45% hydrophobic residues) followed by tandem repeats of Q with either asparagine (Q/N) or histidine (Q/H) followed by a 7 amino acid Q repeat (Fig. 1A). QQ repeats are also observed throughout the RGO protein sequence. *RGO* has been erroneously annotated as repressor *ROX1-like* in TAIR (The Arabidopsis Information Resource) database; *ROX1* encodes a DNA-binding protein that represses the expression of hypoxia genes in yeast (Balasubramanian et al., 1993; Deckert et al., 1995). Protein sequence alignments between *RGO* and the yeast repressor ROX1 showed no homology. As well, RGO does not contain a high-mobility group (HMG) DNA binding domain that is characteristic of repressor ROX1 proteins. Similarly, several genes with deduced protein sequence homology to RGO in *Camelina sativa*, *Brassica rapa* and *Brassica napus* (Supplemental Fig. S2) have also been incorrectly annotated as repressor ROX1 proteins. Proteins with homology to RGO have yet to be identified in the sequence databases of plants outside of the *Cruciferae* species.

**Figure 1.**
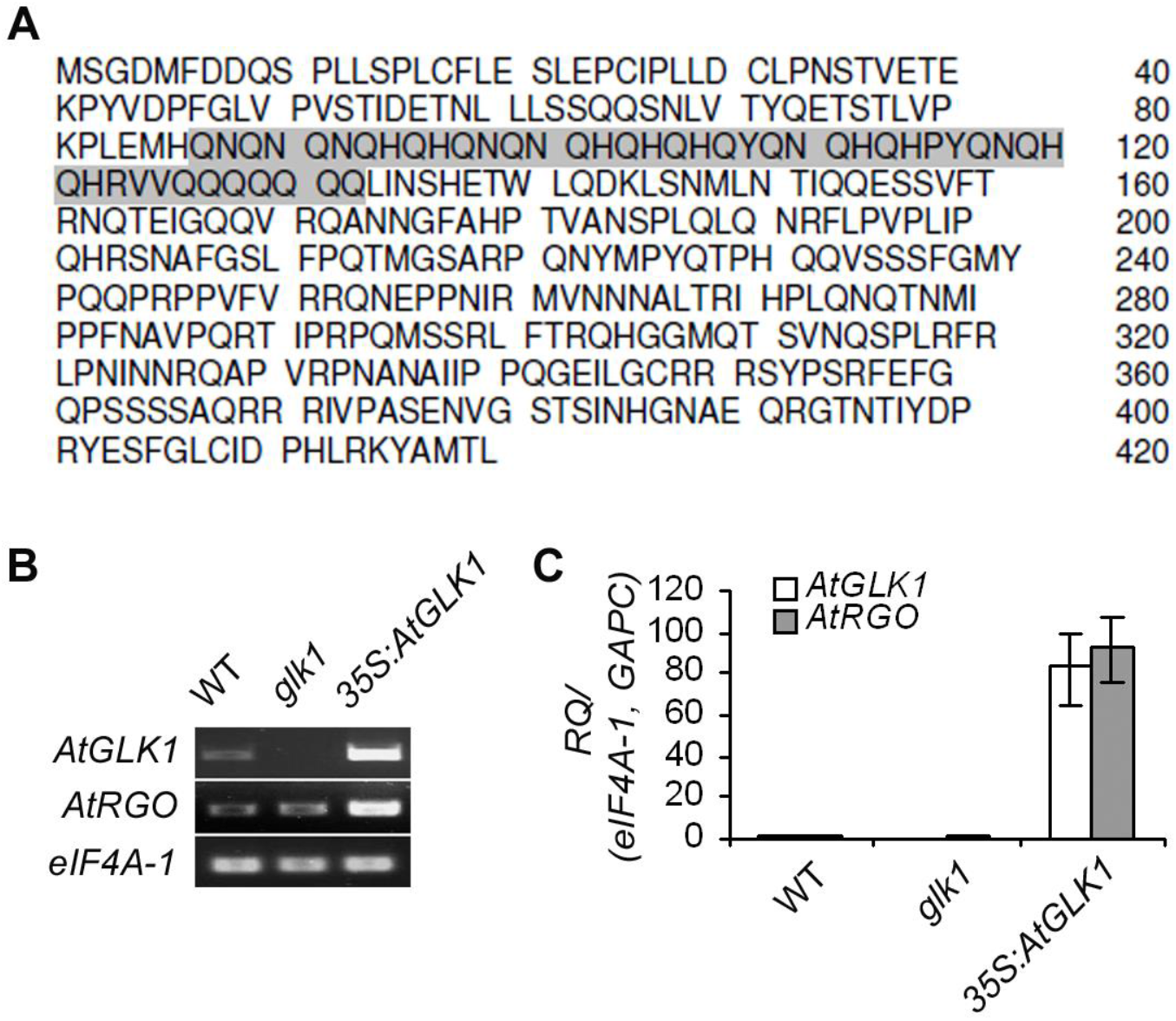
Deduced amino acid sequence and upregulation of *Arabidopsis <u>R</u>esponse to <u>G</u>LK1 <u>O</u>verexpression* (*RGO*, *At1g77960*). (A) amino acid sequence of *At*RGO in single-letter code and the glutamine rich region is highlighted in gray. Numbering of amino acids are on the right. (B) Transcripts accumulation of *AtGLK1* and *AtRGO* in wild-type (WT), *glk1* mutant, and 35S:*AtGLK1* plants with RT-PCR compared to *eIF4A-1*, a housekeeping gene transcripts; (C) *AtGLK1* and *AtRGO* transcripts with qPCR relative to *eIF4A-1* and *GAPC*, two housekeeping gene transcripts. Relative transcripts represent as mean fold change values ± standard error of mean (SEM) from three biological replicates.

### *RGO* expression is ubiquitous in Arabidopsis vegetative and floral tissues

*RGO* transcripts were expressed ubiquitously in Arabidopsis plant tissues such as roots, leaves, stem and internodes, inflorescence apex, and siliques as demonstrated by semi-quantitative RT-PCR (Fig. 2A). *RGO* transcripts were relatively abundant in cauline leaves and scarce in mature siliques as validated by qPCR experiments (Fig. 2B). We also generated transgenic lines expressing a 1.9 kb *RGO* promoter fragment fused to GUS reporter gene in the WT (Col-0) background to localize expression of this gene. Gus expression was most intense in early seed germination, starting one day after germination, and in cotyledons and slowly progressed to vascular tissues in the root (Fig. 2C). In later stages of plant development, expressions were mainly confined to veins of rosette and cauline leaves. In inflorescence tissues, expressions were prominent in the stem, young apex, and young siliques. By contrast, both mature flowers and mature siliques displayed reduced GUS accumulation and no expression was detected in seeds. The *RGO* expression is also diurnally regulated (Supplemental Fig. S3); *RGO* transcripts began to accumulate early in the light cycle and peaked during the middle of the light cycle, then gradually diminished towards the end of the light-cycle and remain unchanged during the dark cycle (Supplemental Fig. S3). This data suggest that *RGO* expression is light or photoperiod sensitive.

**Figure 2.**
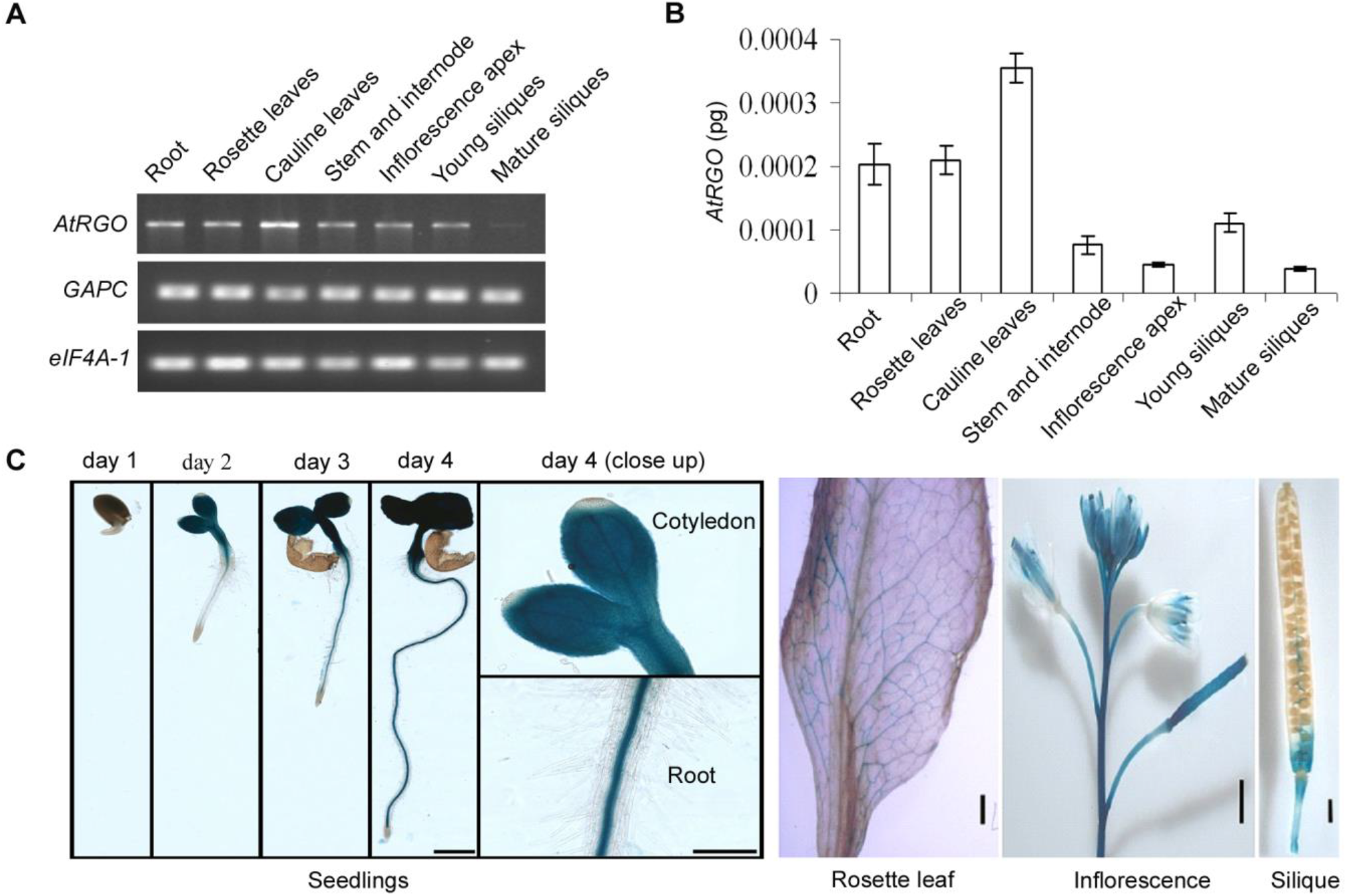
Expression pattern of *AtRGO*. (A) *AtRGO* transcripts accumulation in wild-type (WT) Col-0 with RT-PCR compared to *eIF4A-1* and *GAPC*, two housekeeping control transcripts; (B) Absolute quantification of *AtRGO* transcripts with qPCR. Transcripts represented are mean fold change values ± standard error of mean (SEM) from three biological replicates; (C) *AtRGO* promoter driven reporter GUS reporter gene expression in WT plants. GUS expression was observed starting one day after germination; GUS expression was intense in cotyledons and slowly progressed to vascular tissues in the root. Scale bar, 200 μm in seedlings, and bar, 1 cm in mature tissues.

### RGO is localized to the Endoplasmic Reticulum

The presence of hydrophobic amino acid residues at the N-terminus of the RGO protein suggested that it is most likely membrane localized. As the RGO protein sequence did not reveal any functional domains and organelle localization signals, we undertook cellular localization studies to provide insights into the potential function of RGO. A C-terminal translational fusion of *RGO* with YFP (yellow fluorescent protein) under the control of the Cauliflower Mosaic virus (CaMV) 35S promoter was constructed and stable transgenic Arabidopsis plants were generated. Transgenic plants expressing 35S:YFP were used as a control. As displayed, free YFP fluorescence were observed to be evenly distributed in the cytoplasm and nucleus of leaves of 35S:YFP transgenic plants, (arrows, Fig. 3, A and C). The nuclei were identified by DAPI staining in Fig. 7B. In epidermal cells of leaves expressing 35S:*RGO:*YFP, YFP fluorescence was observed in distinct locations in the cytoplasm and surrounding the nucleus (Fig. 3, D and F). Unlike the control 35S:YFP leaf tissues, no fluorescence was detected in the nucleus (Fig. 3F). The nuclei were identified by DAPI staining in Fig. 7E. This pattern of fluorescence is consistent with proteins localizing to the endoplasmic reticulum (ER) (Nelson et al., 2007; Wulfetange et al., 2011). The presence of the hydrophobic N-terminus of RGO likely enables it to be localized to the membranes. To confirm these findings, FM4-64FX, a plasma membrane specific fluorescent dye was used in combination with YFP fluorescence to demonstrate that 35S:RGO-YFP fluorescence (yellow) was distinctly separated from the FM4-64FX (blue) plasma membrane stain (Fig. 3H), whereas, free fluorescence in the control leaf tissues (35S:YFP) overlapped (white colour) with the plasma membrane as well as the nucleus (Fig. 3G). These fluorescence patterns were indicative of ER localization and suggest that RGO is associated with the ER and not the plasma membrane. To further confirm the ER localization of RGO protein, we compared transient overexpression of an ER marker, 35S:GFP-HDEL (Batoko et al., 2000) to the expression of 35S:RGO-YFP in tobacco leaves. The expression pattern of RGO (Fig. 3I) is identical to the ER expression pattern of 35S:GFP-HDEL (Fig. 3J) and confirms that RGO is localized to the ER.

**Fig. 3.**
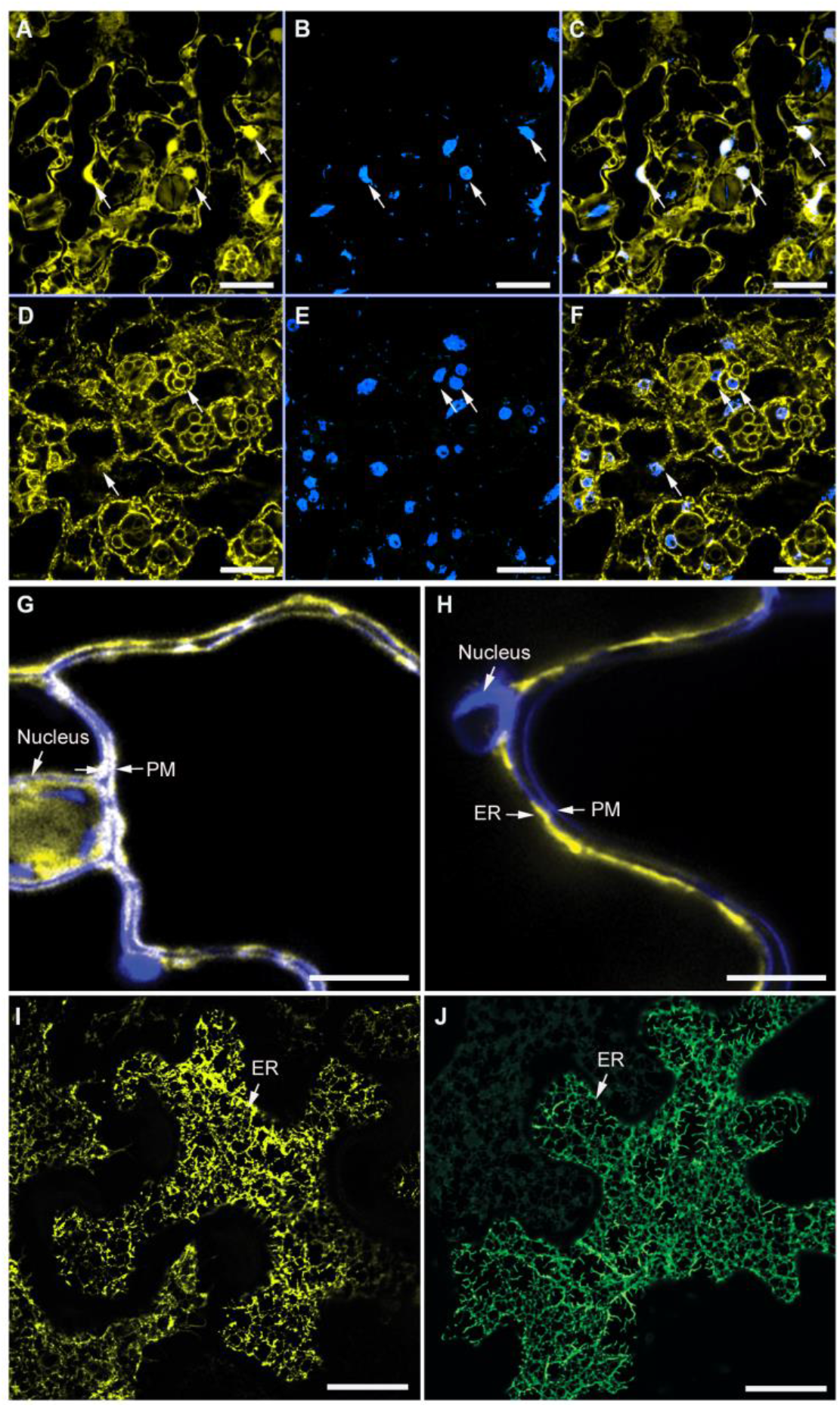
Subcellular localization of *At*RGO-YFP fusion protein by confocal microscopy in transgenic Arabidopsis leaves and tobacco leaves. Plants expressing 35S:YFP were used as positive control (A-C) for comparing *At*RGO-YFP fusion expression (D-F). (A and D), YFP signal in yellow; (B and E), DAPI strained nuclei in blue; (C and F) merged. Arrow head shows the nucleus and endoplasmic reticulum (ER). Zoom in view of 35S:YFP (G) and *At*RGO:YFP (H) with FM4-64FX stain. YFP fluorescence in yellow and plasma membrane stain in blue, a false colour for contrast is shown in G-H and arrow head shows the nucleus and plasma membrane (PM) in G-H. Transient expression of *At*RGO-YFP (I) and GFP:HDEL, an ER marker (J) in tobacco leaves. Arrow head shows the ER in I-J. Scale bar, 20 μm in A-F and 5 μm in G-J.

### *RGO* is required for root development, normal growth and flowering

Phenotypic studies were undertaken to further elucidate the function of *RGO*. We obtained three T-DNA insertion lines (Salk_025545, Salk_109614, and Salk_109424) for *RGO* from the Arabidopsis Biological Resource Center (ABRC), Ohio State University, USA. Arabidopsis T-DNA design tool (http://signal.salk.edu/tdnaprimers.2.html) indicated that Salk-109424 has a second T-DNA insertion in gene *At5g33175*. We could not detect T-DNA insertions in the *RGO* gene in the Salk_109424 line (data not shown). However, we identified a T-DNA insertion in the first exon of *RGO* at 33 bp down stream of the transcription start site in the Salk_109614 plants (Supplemental Fig. S4A). Similarly, we identified a T-DNA insertion in Salk_025545 in the fifth intron at 1170 bp down stream of transcription start site (Supplemental Fig. S4A). Both Salk_109614 and Salk_025545 T-DNA insertion lines produced uninterrupted full length *RGO* transcripts (Supplemental Fig. S4D), which was verified by sequencing the *RGO* gene transcripts from both Salk_109614 and Salk_109424 plants and showed that the transcripts were identical to the WT.

As we could not isolate homozygous T-DNA insertional knockout lines for *RGO*, we generated artificial micro RNA (amiR) silencing lines for the gene. We screened 72 Basta resistant primary transformed plants and identified two lines (*rgo*-1 and *rgo*-2) and transcripts analyses of T2 generations of by RT-PCR as well as qPCR revealed that these two lines had significant reductions in transcript accumulation of *RGO* (Fig. 4 A). Plants from both *rgo* lines exhibited a minimum of two-week delay in bolting compared to the WT (Fig. 4, B and C). These plants were able to set seeds albeit to much lesser degrees than WT plants and with the phenotypes being stable in subsequent generations. Seeds from subsequent generations (T4) showed substantial decreased germination rates (70-80%) compared to the WT (Supplemental Fig. S5A). To validate the phenotypes observed in the *RGO* silenced (*rgo*) lines, we generated Arabidopsis lines that constitutively expressed *RGO* (35S:*RGO*). Section and staining of *rgo* seeds showed impaired embryo development compared to the WT and to the 35S:*RGO* plants. In the *rgo* lines, the majority of embryos were immature, which corroborated with the lethality of seeds and consequently decreases in seed germination. (Supplemental Fig. S5B). In light of these findings that linked *RGO* with plant development in vegetative tissues, we examined the effect of *RGO* silencing on root growth. Compared to the WT seedlings, *rgo* seedlings grew significantly slower and with a 67% decrease in root length compared to the WT, whereas the root length of 35S:*RGO* seedlings were similar to the WT (Fig. 5, A and B). Collectively, these results suggest that *RGO* is required for normal growth as well as viable seed development.

**Figure 4.**
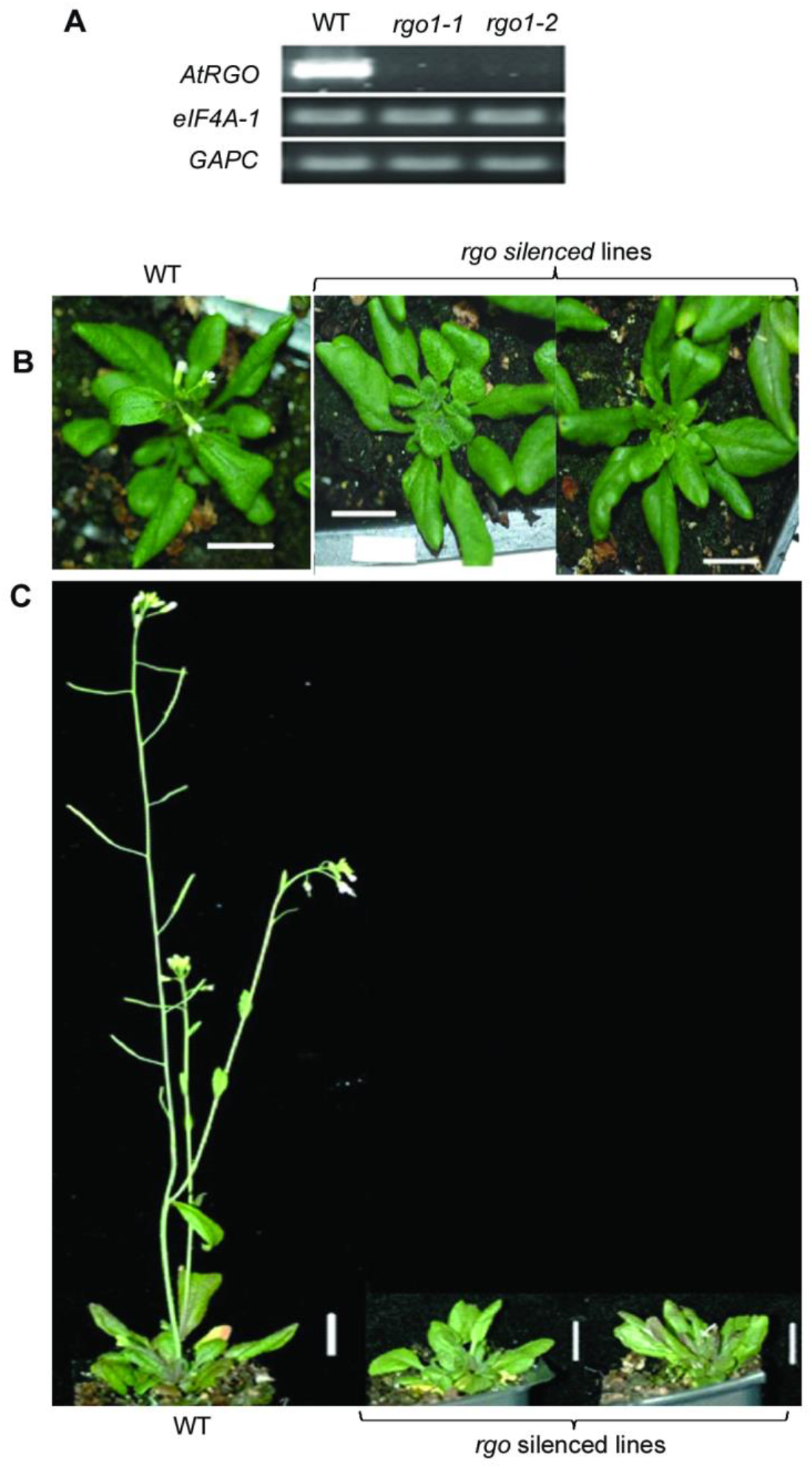
Phenotype of *rgo* plants compared to WT (Col-0). (A) A RT-PCR of full length *AtRGO* transcripts in WT and *rgo* plants compared to *eIF4A-1* and *GAPC* transcripts; (B) qPCR of *AtRGO* transcripts in WT and *rgo* plants; (C) phenotype at four-week age; (D) phenotype at six-week age. Scale bar, 1 cm in C-D.

**Figure 5.**
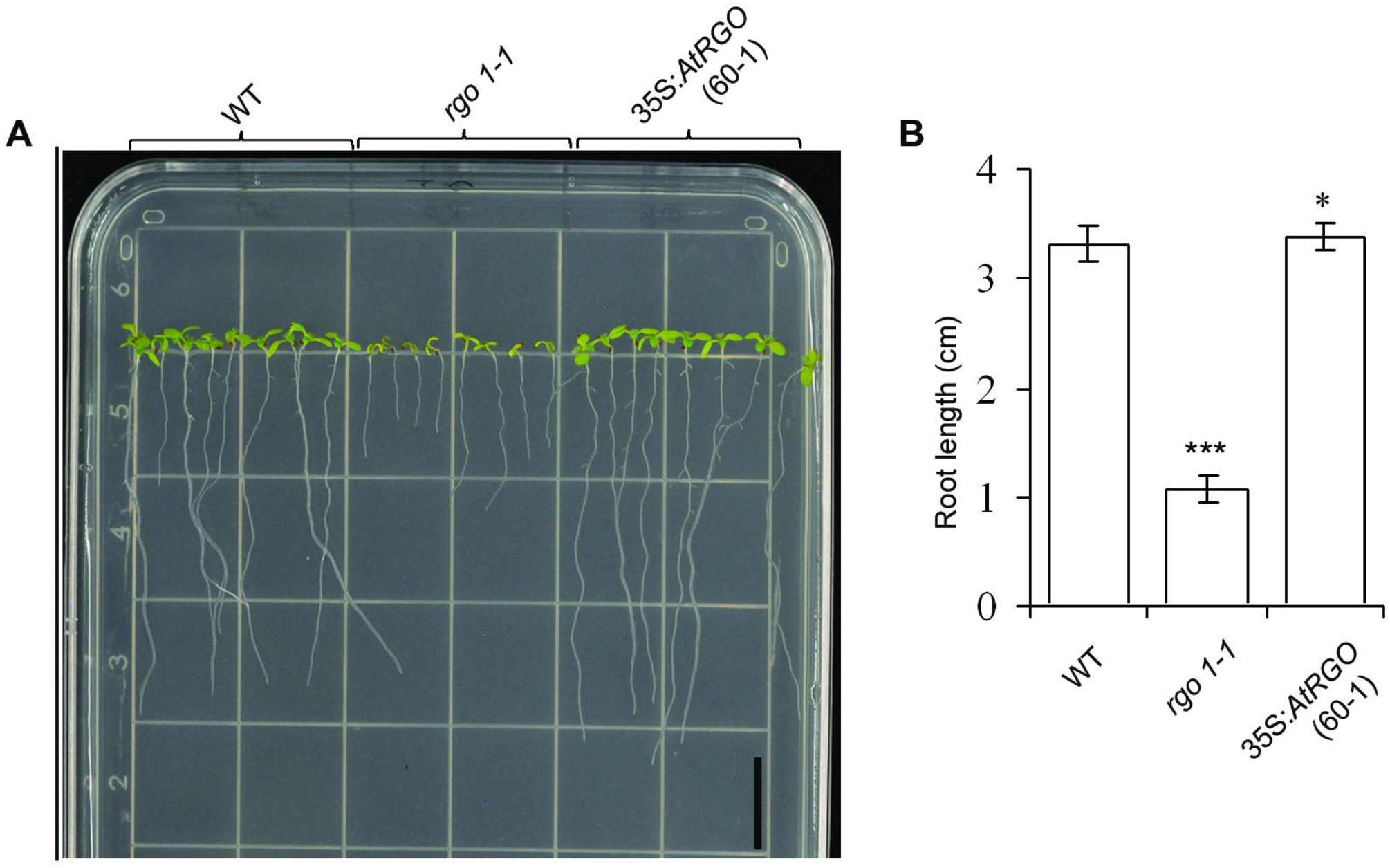
Root growth phenotype of WT, *rgo* and *35S: AtRGO* lines at seven days post germination. (A) representative images on minimal media (MM), (B) corresponding root length. Error bar = standard error of mean. Scale bar, 1 cm in A. Statistical significance of root lengths were analyzed using student’s t-test: Two sample assuming unequal variance at 95% confidence level. *** represents the p value < 0.00000001, and * represents p value > 0.1.

### Constitutive expression of *RGO* increases growth and seed yield

Analyses of T3 generations of three independent 35S:*RGO* lines indicated a significant accumulation of *RGO* transcripts, with line 60-1 exhibiting a 20-fold increase in transcript accumulation compared to the WT (Fig. 6, A and B). All the lines overexpressing *RGO* showed an acceleration of flowering by one-week, when compared to the WT under long-day growth conditions (Fig. 6C). Additionally, aerial vegetative growth were observed 2-3 weeks earlier compared to the WT and by week four, the transgenic plants showed increased numbers of axillary vegetative buds per rosette compared to the WT resulting in increased number of shoots per rosette (Fig. 6D). We also observed that the length of the rosette leaves in *RGO* transgenic plants were considerably longer than that of the WT (Supplemental Fig. S6). Given the significant increase in biomass and shoots, we assessed seed yield in these transgenic plants overexpressing *RGO*. On average, there was a 20 % higher seed yield than the WT under identical growth conditions (Fig. 6E). We were interested to know if the seed yield phenotype can be recapitulated in related Brassica species. We generated two lines of *Brassica napus* overexpressing *AtRGO* (35S:*RGO*:). Stable transgenic *B. napus* (T3) lines overexpressing *RGO* showed similar increased vegetative branching compared to the WT (Supplemental Fig. S7) with an average 18 % increase in seed yields (Fig. 6F) when grown under controlled environmental conditions.

**Figure 6.**
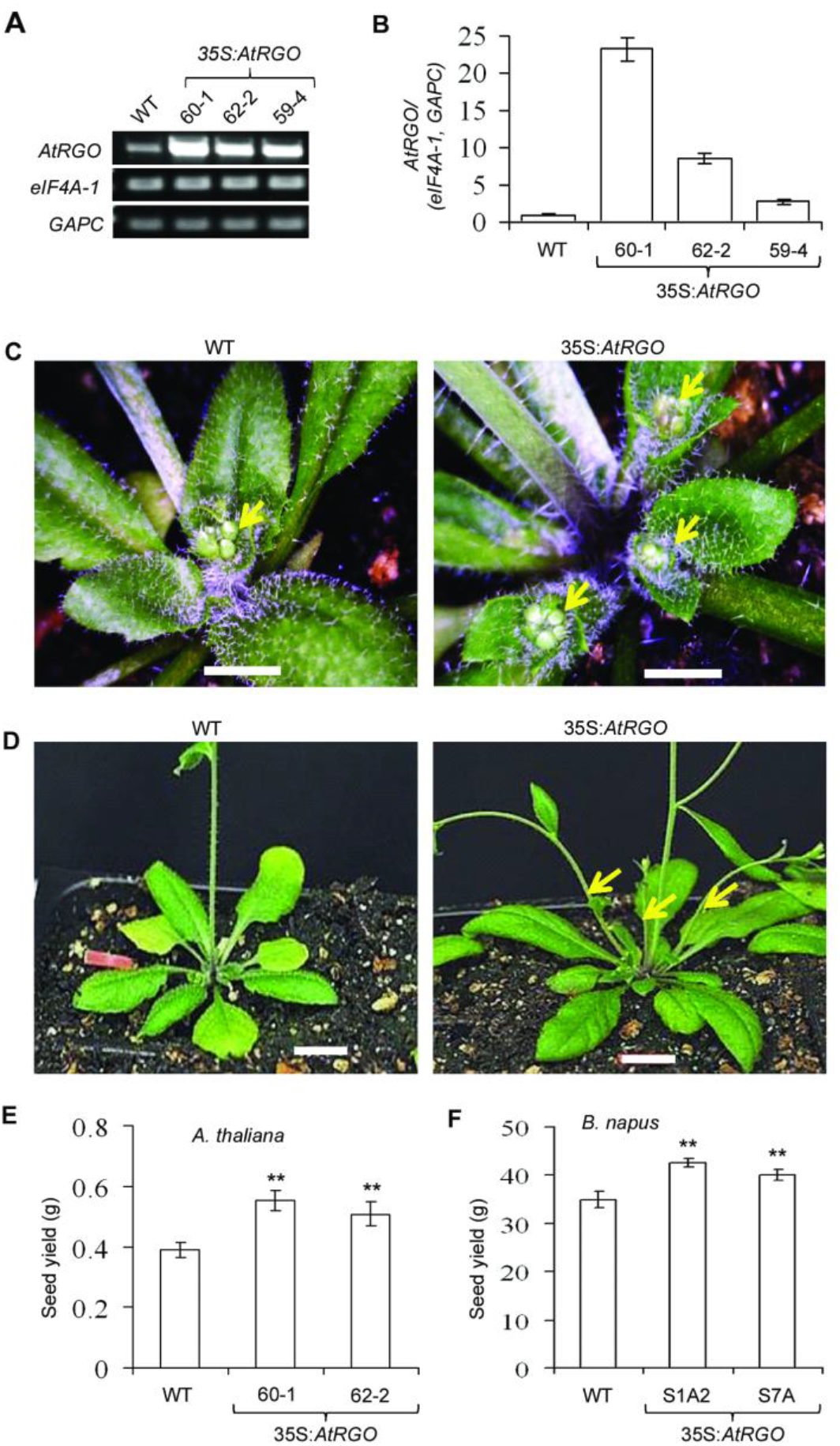
Phenotype of *35S:AtRGO* plants compared to WT. (A) A RT-PCR of full length *AtRGO* transcripts compared to *eIF4A-1* and *GAPC* transcripts in WT and three independent lines of *35S:AtRGO* plants; (B) qPCR of *AtRGO* transcripts in WT and *AtRGO* overexpressing lines; (C) Four-week old plants; (D) Five-week old plants. Emergence of inflorescence buds are shown with arrow head in C and axillary buds in D; (E) Average seed yield from 10 Arabidopsis plants of WT and *35S:AtRGO* from a long-day photoperiod growth conditions; (F) seed yield from individual plants of *B. napus* in WT and *35S:AtRGO* (two lines). Scale bar, 1cm in C-D. Statistical significance was analysed using student’s t-test: Two-sample assuming unequal variances at 95% confidence level compared with empty vector. The p value for each analysis is shown as asterisk, where p value ≤ 0.001 (**). Error bar = standard error of mean.

### *RGO* mediates CK responses to regulate endogenous CK levels and safeguard the positive effects of CK in plant development

To explore if a relationship exists between *GLK1* overexpression, *RGO* upregulation and CK responsiveness, we first examined if ectopic overexpression of *GLK1* has an effect on sensitivity to CK signalling using the green calli/shoot regeneration assay (Kiba et al., 2004). Robust green calli formation was observed for external application of 0.005 μM BA for *GLK1* overexpressing plants compared to 0.1 μM BA for the WT (Fig.7A). This observation prompted us to use *RGO* silencing and overexpression to determine if RGO can, in part, be a factor in CK response. One of the well-documented effects of exogenous CK on plants is shortened root growth (Skoog and Miller, 1957) and that CK levels regulate root development (Werner et al., 2001; Werner and Schmülling, 2009). As observed, *rgo* lines display shorter root growth compared to the WT and *RGO* overexpressor seedlings (Fig. 5A). Therefore, we compared the root growths of the WT, *35S:AtRGO* and *rgo* seedlings in the presence of 5 μm Zeatin, a synthetic CK. The effect on root growth inhibition by Zeatin was partially mitigated in *RGO* overexpressing seedlings compared to the WT and *rgo* seedlings suggesting that overexpression of *RGO* can potentially dampen the inhibitory effects of CK on root development (Fig. 7B). In addition to the effects on root growth, it is known that external applications of CK also facilitate green calli and shoot regeneration. Green calli formation from excised hypocotyl was evaluated to measure CK responses (Kiba et al., 2005). The results showed that while the *RGO* overexpressing and WT plants showed minimal differences in their response to exogenous applications of BA, the *rgo* plants were unable to respond to BA and did not induce green calli formation (Fig. 7B). External applications of CK has also been shown to improve chlorophyll retention and delay dark-induced senescence in leaves (Gan and Amasino 1995; Kim et al 2006; Talla et al 2016). As well, increased endogenous levels of CKs have been shown to be able to delay senescence in plants (Richmond and Lang, 1957; Lin et al., 2002; Ma and Liu, 2009; Zhang et al., 2010; Liu et al., 2012). We assessed chlorophyll retention in detached leaves after dark-induced senescence in the absence or presence of BA. After seven days of dark incubation in the absence of BA (7d-BA), neither the WT nor the *rgo* plants retained any chlorophyll (Chl) (Fig. 7D). In contrast, 35S:*AtRGO* leaves retained 26% of the Chl content (Fig. 7D). In the presence of BA (7d +BA), 35S:*AtRGO* plants did not lose any Chl (Fig. 7D). Interestingly, in the 7d+BA treatment, the *rgo1-1* silenced line also retained significantly more Chl than the WT (Fig. 7D). This retention could result from increased CK accumulation in the silenced line facilitating some signalling even though *RGO* is knocked down. Analyses of bioactive CK levels confirmed that that both trans-zeatin (tZ) and 2-isopentenyladenine (2iP) were significantly elevated in the *rgo* plants compared to the WT (Fig. 7E). There was a modest but significant decrease in 2iP levels in *RGO* overexpressing plants (Fig. 7E). This increase in endogenous CK levels in *rgo* plants (Fig. 7E) is reflected in lower transcript levels encoding CK degrading enzymes CK oxidases *CKX4* and *5*, (Fig. 7F). Taken together, these results suggest that *RGO* can respond to modulate CK levels in a fashion that facilitates or promotes favorable plant processes. It can attenuate the negative effect of external CK application on root growth and conversely, positively facilitate CK induced delay of senescence in excised leaves.

**Figure 7.**
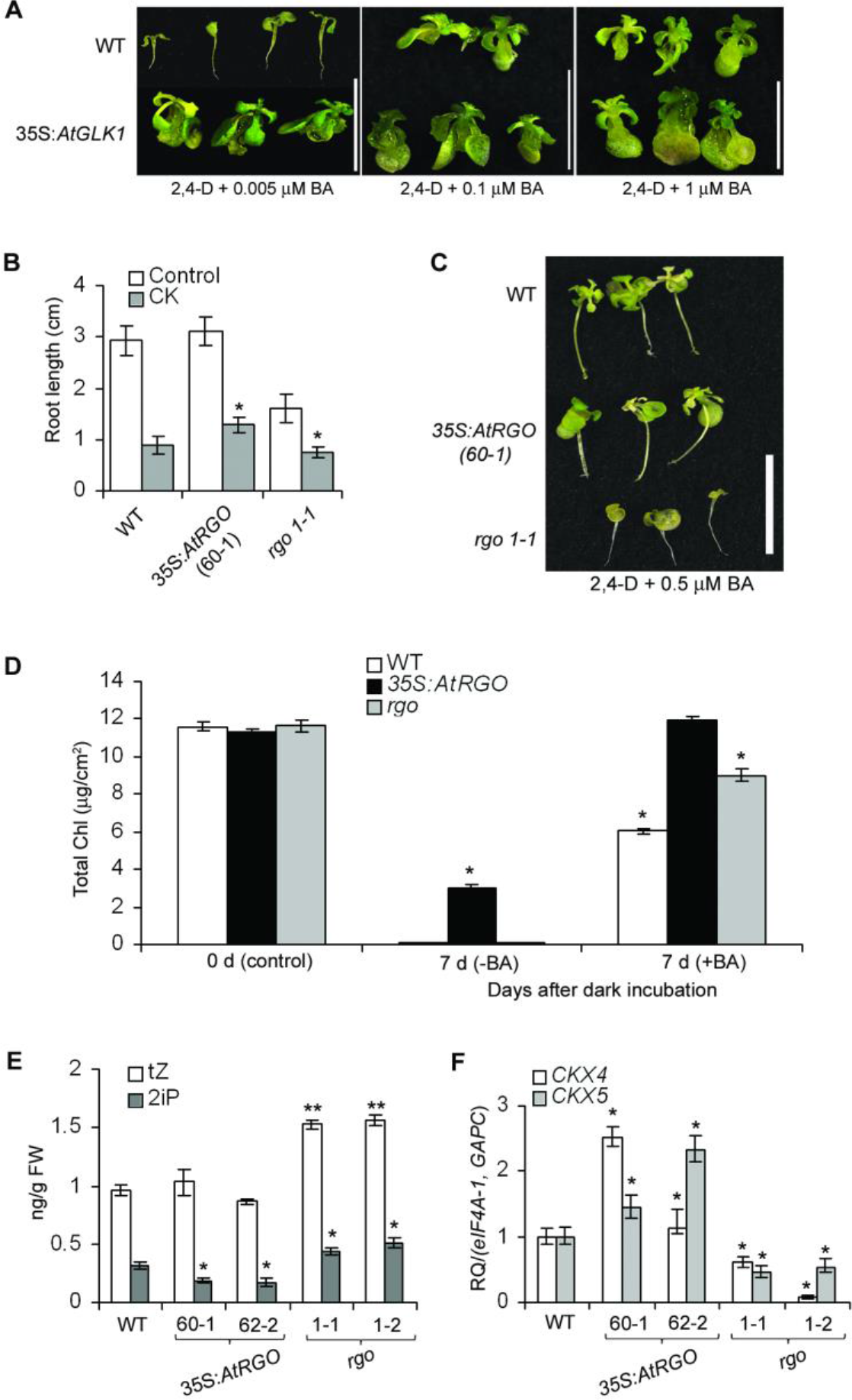
Cytokinin (CK) response, CK levels in 35S:*AtRGO* and *rgo* silenced plants. (A) Hypersensitive response of *35S:AtGLK1* plants to BA, a synthetic cytokinin (CK), in green calli formation. WT and *35S:AtGLK1* seeds were germinated in dark for five days on half strength MS agar plates then elongated hypocotyls were excised out and grown on half strength MS agar plates supplemented with 0.005 μM 2,4-D plus varying concentration (0.005 to 1 μM) of BA under long-day photoperiod conditions. After 30 days representative callus were photographed. Scale bar = 1 cm. (B) Root length of seedlings at seven days post germination on minimal agar media in absence or presence of 5 μM CK, *represents the p value < 0.001; (C) Green callus formation from excised hypocotyl in presence of 0.005 μM 2,4-D plus 0.5 μM BA (CK) on MS agar plates of WT, 35S:*AtRGO* and *rgo*. Scale bar = 1 cm. (D) Effect of exogenous application of CK on leaf chlorophyll (Chl) retention during dark-induced senescence; (E) Bioactive CK [zeatin (tZ, open bar) and 2-isopentenyladenine (2iP, grey bar)] level in four-week old rosette leaves. p value < 0.001 (**) and p value <0.01 (*). Error bars = SE (n=3). * denotes the significant difference between −BA and +BA treatment (p<0.001, Student’s t-test); (F) Relative transcripts of *CKX4* and *CKX5* quantified with qPCR in four week old leaves of wild-type (WT) Col-0, 35S:*AtRGO*, and *rgo*, compared to *eIF4A-1* and *GAPC*, two housekeeping control transcripts. Relative transcripts (RQ) represent as mean fold change values ± standard error of mean (SEM) from three biological replicates. Error bars = SE (n=3). * denotes the statistical significant (p<0.001, Student’s t-test).

## Discussion

*RGO* (*At1g77960*) is a single copy gene in Arabidopsis and encodes a protein of unknown function. *RGO* transcript levels increased highly in Arabidopsis plants ectopically overexpressing either the Arabidopsis *GLK1* or wheat *TaGLK1* transcription factor. *RGO* expression however did not seem to be regulated directly by *GLK1* as *glk1* plants did not show any down regulation of *RGO*. To explore if *RGO* expression was related to requirement for CK levels rather than directly to GLK1 overexpression, we measured *GLK1* and *RGO* transcript levels in *atipt1 3 5 7*, a quadruple CK biosynthetic isopentyltransferases mutant severely deficient in endogenous iP- and *t*Z-type cytokinins (Miyawaki et al., 2006). Interestingly, *GLK1* levels were increased over 30-fold in this mutant, but increases in *RGO* levels were not observed (Supplemental Fig.S8). This indicated that increased transcript levels of *RGO* and *GLK1* does not necessarily occur in tandem and suggests that *GLK1* does not directly regulate *RGO*. It is not known why *GLK1* is so highly upregulated in this mutant. It is well established, however, that *GLK1* gene transcription and protein accumulation is regulated by plastid retrograde signals (Martin et al., 2016; Tokumaru et al., 2017) and it is possible to consider that endogenous CK levels may have an effect on this signalling. Nevertheless, this result further underscores the requirement of *RGO* in CK response. The genes directly regulating transcription of *RGO* is not yet known. The elucidation of the transcriptional elements directly regulating the expression of *RGO* will have to await identification of binding domains in the promoter region of *RGO* to enable yeast-one-hybrid screens. Unlike *GLK*, *RGO* is not directly involved in the regulation of chloroplast development as *RGO* silenced plants did not show a pale green phenotype (Fig. 4, C and D) observed in the *glk1glk2* plants, which are impaired in chloroplast development (Yasumura et al., 2005; Waters et al., 2008). Moreover, expression of *RGO* (Fig. 3) in all tissue types including photosynthetic tissues also argues against a role critical to chloroplast development. Ectopic overexpression of *RGO* in the pale green *glk1 glk2* background did not rescue chloroplast development, suggesting that it does not act upstream of the *GLKs* (unpublished). More likely, upregulation of *RGO* expression is a response to ectopic overexpression of *GLK1* or type-B ARRs without a phosphor-relay regulatory capacity such as *ARR21C* (Tajima et al., 2004; Kiba et al., 2005) in plants with normal endogenous CK levels.

### *RGO* is essential for normal plant development

*RGO* is a single copy gene and the inability to acquire homozygous *RGO* knockout lines without *RGO* transcripts is suggestive that it is essential in plant development. We used amiR silencing as well as ectopic overexpression of *RGO* in Arabidopsis to gain insights into its role in plant development. Observations of reduced transcripts in the *RGO* downregulated (*rgo*) plants were correlated to displays of alterations in normal vegetative and root developments including growth retardation, low seed yields and seed quality leading to lower germination rates in successive generations (Fig. 4, Fig. 5, Supplemental Fig. S5). Conversely, Arabidopsis plants ectopically overexpressing *RGO* consistently produced higher seed yields and vegetative shoots.

We hypothesized that at least one explanation for the phenotypes produced by *RGO* overexpression and *RGO* knockdowns could be the result of the modulations of CK response by *RGO*. To this end, we carried out experiments to test specifically whether *RGO* is involved in attenuating or facilitating CK responses. We provide several lines of evidence to support this hypothesis. Firstly, overexpression of *RGO* dampened the effects of external application of CK on root development (Fig. 7A). Conversely, *RGO* silenced (*rgo*) plants showed retarded root growth even in the absence of external CK application (Fig. 5, A and B). Secondly, excised hypocotyls from *RGO* silenced plants were unable to produce green calli/shoot formation in external application of CK (Fig. 7B). Thirdly, increased expression of CK oxidase genes, *CKX4* and *CKX5* in leaves of *RGO* overexpressing plants (Fig. 7E) suggested that one role of the involvement of *RGO* in CK response is to monitor and respond to endogenous levels of CK through *CKX* expression. Elevated CK levels can be attributed to modulation of *CKX* that will prevent the degradation of CKs (Mok and Mok, 2001; Werner et al., 2001; 2003). Conversely, knockdown of *RGO* produced significant increases in bioactive CK concomitant with significant decreases of *CKX4* and *CKX5* expression in leaves of the *rgo* plants (Fig. 7D,E). *AtRGO* was identified as a candidate in a GWAS (<u>g</u>enome <u>w</u>ide <u>a</u>ssociation <u>s</u>tudy) study to elucidate genes in shade avoidance (https://explorations.ucdavis.edu/docs/2015/choi.pdf). There was however, no indication the report was peer reviewed and that whether the knockouts of *AtRGO* were genotyped to assure homozygosity as well as absence of *AtRGO* gene transcripts. Nevertheless, it is known that shade avoidance is in part hormonally regulated and involves auxin and cytokinin levels mediated by leaf CKXs (Wu et al., 2017; Yang and Li, 2017). It is conceivable therefore that *AtRGO* could play a role in shade avoidance and should offer opportunities for future studies.

### Modulation of CK response by *RGO* leads to increased seed yield

Ectopic overexpression of *RGO* significantly increased seed yield in both Arabidopsis and *B. napus*. It was observed that the Arabidopsis *ckx3 ckx5* double mutant had increased CK levels resulting in increased floral meristems leading to increased seed yield (Bartrina et al., 2011). It is known that tissue specific expression of *CKXs* influence CK levels in those tissues (Mok and Mok, 2001; Werner et al., 2001; 2003). Similarly, silencing of *OSCKX2* in rice increased CK accumulation in inflorescence meristems and increased tiller number resulting in enhanced grain yield (Ashikari et al., 2005; Yeh et al., 2015). Grain weight in wheat is also associated with *TaCKX6-*D1, an orthologue of *OSCKX2* (Zhang et al., 2012). Silencing of barley *HvCKX* genes has also been observed to increase seed yield (Zalewiski et al., 2010, 2012, 2014). It is conceivable that increased seed yields in *RGO* overexpressing plants are, in part, a result of enhanced CK signaling. The reduction of *CKX3* and *CKX5* transcript levels in inflorescence buds of *RGO* overexpressing plants (Supplemental Table S4) supports the view that enhanced effects of CK responses are influenced by *RGO*.

The mechanism by which *RGO* facilitates CK responses or regulate plant development remains to be elucidated. *RGO* encodes a glutamine (Q) rich protein containing domains of Qs and tandem QNs or QHs. Other than these features, and a hydrophobic N-terminal, RGO contains no known identifiable protein domains. The absence of localization of RGO to the nucleus (Fig. 3) precludes direct functional interactions with transcription factors. Proteins with high QN contents and poly-Q stretches have been associated with neurodegenerative disorders in humans through protein aggregation (Perutz, 1994; Guo et al., 2007; Kuiper et al., 2017) and expansion of Q stretches has been suggested to even modulate the aggregative properties of flanking amino acids (Kuiper et al., 2017). Little is known however of the functional roles of high Q containing proteins in plant development, although Q and HQ domains have been implicated in protein binding/aggregation (Guan et al., 2017) and in RNA binding (Muthuramalingam et al., 2016) in plants. As both poly Q and poly HQ domains are prominent in the RGO protein (Fig. 1A) and as the RGO protein is localized to the ER., we speculate that RGO may interact with CK signaling components as CK signaling components reside in the ER (Caesar et al., 2011; Lomin et al., 2011; Wulfetange et al., 2011; Lomin et al., 2018; Romanov et al., 2018) as well as mediating CK homeostasis by CKXs which takes place in the ER (Schmülling et al., 2003; Werner et al., 2003). Mechanistic aspect of RGO at protein level is under investigation. Identification of transcription factors that directly regulate *RGO* transcription as well as RGO protein/protein interaction partners will aid in elucidation of a mechanism of RGO in plant development.

## Materials and Methods

### Plant material

For routine propagation of *Arabidopsis thaliana*, seeds from wild-type (Col-0), T-DNA lines (Salk-025545, Salk-109614, Salk-109424), *RGO* silenced lines, 35S:*RGO, glk1* and the *glk1 glk2* double-knockout lines (N9806, N9807, respectively, obtained from the Nottingham Arabidopsis Stock Centre (NASC), Nottingham, UK.), the 35S:*GLK1* and 35S:*TaGLK1* lines were surface sterilized with 2% bleach (v/v) plus 0.001% tween-20, washed five times with sterile water and stratified for two days at 4°C. Seeds were grown in 48 cell trays on sterilised PRO-MIX MPV potting mixture (Premier Tech Horticulture, Rivière-du-Loup, QC, Canada) and grown in a Conviron PGC 20 CMP model 6050 cabinets (Winnipeg, Manitoba, Canada) at 21-22 °C, 40-60 % humidity, under long-day conditions (16-h-light/8-h-dark cycle) and a fluorescent light intensity of 100-120 μmol photons m^−2^ s^−1^. Plants were fertilized once a week using 1g per litre solution of 20-20-20 fertilizer (Master Plant-Prod Inc. Brampton, ON, Canada). Similarly, sterilized seeds of *Brassica napus cv* Westar (wild-type) or *RGO* overexpressing transgenic plants were grown in six inch round pots with sterilized mixture of 50% Promix, 25% soil and 25% sand in a Conviron as described above under long-day conditions with a light intensity of 750 μmol photons m^−2^ s^−1^. A solution of two grams per litre 20-20-20 fertilizer was given once every third day for the first two weeks and then once a week until seed set. Ten plants each from *Brassica napus* wild-type and two-independent transgenic lines were grown in growth cabinets with identical growth conditions. Seeds were collected from each individual plant and weighed.

### Promoter GUS construct and analyses

A 1.9 kb upstream of the transcription start site ATG of the *At1g77960* gene was PCR amplified with RGO-promo-F and RGO-Promo-R primer pairs using Phusion® High-Fidelity DNA Polymerase (New England Biolabs, Whitby, ON, Canada) with Arabidopsis genomic DNA as the template and was verified by DNA sequencing (Eurofins Genomics, Louisville, KY, USA). The resulting PCR fragment was gel purified and cloned into the gateway entry vector pENTR®/D-TOPO® (Thermo Fisher Scientific, Waltham, MA, USA). The *RGO* promoter: pENTR®/D-TOPO® plasmid was transferred to the binary vector, pMDC162 (Curtis and Grossniklaus, 2003) using LR clonase II (Thermo Fisher Scientific, Waltham, MA, USA) to generate transcriptional fusion with GUS reporter gene. *Agrobacterium tumefaciens* strain GV3101 -pMP90RK containing the *RGO* promoter: pMDC162 plasmid was transformed to Arabidopsis wild-type (Col-0) plants by floral dipping (Clough and Bent, 1998). Primary transgenic plants (T_0_) were selected on Murashige and Skoog (MS) basal media plus agar plates in presence of 25 μg ml^−1^ hygromycin. At least 10 independent transformed lines from T1 through T3 generation were chosen for GUS analyses. GUS staining was performed as previously described (Murmu et al., 2010). Representative images from three independent lines were presented in figure. All the cloning primers are found in Supplemental Table S2.

### Constructs for subcellular localization of full length *RGO*

An overlap extension approach was used to create a C-terminal translational fusion of RGO with YFP. The *RGO* coding sequence (CDS) without the stop codon was PCR amplified from wild-type (WT) leaf cDNA with RGO-cDNA-F plus YFP-RGO cDNA-R primer pairs that generated a 1284 bp PCR product with an overlap of 20 bp with the N-terminal of YFP. Next, *YFP* was PCR amplified using RGO-YFP-F plus YFP-R primer pairs that generated a 743 bp PCR product with an overlap of 20 bp at the C-terminal of *RGO* CDS. The above two-PCR fragments were then used as templates for PCR amplification with RGO-cDNA-F plus YFP-R primer pairs to generate a 1987 bp *RGO-YFP* translational fusion construct. The resulting PCR product was cloned into the Gateway entry vector pENTR/D-TOPO. Similarly, the YFP CDS was PCR amplified using YFP-F4-Topo plus YFP-R primer pairs and cloned into pENTR/D-TOPO for positive control. All constructs were verified by DNA sequencing. The *RGO-YFP* construct was then transferred to the pK7GW2D binary vector downstream of the Cauliflower Mosaic Virus (CaMV) 35S promoter (35S), using LR clonase II (Thermo Fisher Scientific, Waltham, MA, USA). Similarly, the YFP construct was transferred to the pMDC32 binary vector downstream of the CaMV 35S promoter (Curtis and Grossniklaus, 2003). The resulting 35S:*RGO-YFP*:pK7GW2D and 35S:YFP:pMDC32 plasmids were introduced into *Agrobacterium tumefaciens* and were used to transform WT (Col-0) Arabidopsis plants by floral dipping as described in the previous section. Primary transgenic plants were selected on MS agar plates in the presence of 25 μg ml^−1^ hygromycin for the construct in pMDC32 or in the presence of 50 μg ml^−1^ kanamycin for the construct in pK7GW2D.

### Construction of *RGO* for overexpression studies

A full length CDS of *RGO* was PCR amplified from WT leaf cDNA with Xba1-*RGO*-F and Kpn1-*RGO*-R primer pairs. The resulting 1.3 kb DNA fragment was fused to the CaMV 35S promoter at the Xba1 and Kpn1 sites in the pHS723 binary vector that also constitutively express the GUS reporter gene (Nair et al., 2000). The 35S:*RGO*:pHS723 plasmid was transformed into *A. tumefaciens* as described in the previous section. Primary transgenic plants were selected on MS agar plates in the presence of 50 μg/ml kanamycin and highly expressing transgenic plants were further confirmed by GUS staining. *B. napus* cv Westar transformation with the 35S:*RGO*:pHS723 construct was carried out as previously described (Savitch et al., 2005).

### Construction of artificial microRNA (amiR) to silence *RGO*

Artificial microRNA (amiRNA) gene silencing has been efficiently characterized in *Arabidopsis* (Ossowski et al., 2008). The detail of primer designing and step wise procedure for generating amiRNA constructs can be found in Web MicroRNA Designer (http://wmd3.weigelworld.org). The Web MicroRNA Designer was used to generate a 21mer amiRNA (TAACGGAATACACGGTTGCGG) sequence found within the CDS of *RGO* between 68-84 bp to silence *RGO*. Based on this 21mer amiRNA sequence, four primers: [I: microRNA forward (I miR-S68), II: microRNA reverse (II miR-A68), III: microRNA* forward (III miR* S68), IV: microRNA* reverse (IV mir*A68)] were used to engineer the 21mer amiRNA by site-directed mutagenesis into the Arabidopsis endogenous plant microRNA, miR319a. The engineered miRNA319a targets the *RGO* gene for silencing. The following steps were used to engineer the amiRNA and amiRNA* sequences into an endogenous miR319a precursor. First, three PCR fragments using the pRS300 plasmid as template that harbour the miRNA19a were generated. The first PCR amplified a 271 bp fragment with pRS300 A plus IV mir*A68 primer set that generated overlap with one end of the pRS300 plasmid and mir*A68. The second PCR amplified a 170 bp product with III miR* S68 plus II miR-A68 primer set that generated overlap with mir*A68 and miR-A68. The third PCR amplified a 290 bp product with I miR-S68 plus pRS300 B primer set that generated overlap with miR-A68 and other end of pRS300 plasmid. Finally, an overlap PCR was performed using the above three PCR fragments as templates with pRS300 A plus pRS300 B primer set that generated a 699 bp fragment of engineered miRNA19a with overlap of pRS300 destined for subsequent cloning. The engineered miRNA19a was cloned into pJET1.2 (Thermo Fisher Scientific, Waltham, MA, USA) and verified by sequencing (Eurofins Genomics, Louisville, KY, USA). The engineered miRNA19a was excised from pJET1.2 with BamH1 and EcoR1, and ligated into the binary vector pBAR1 under the CaMV 35S promoter (35S:amiRNA19a:pBAR1). The construct was transformed to Arabidopsis WT plants via *A. tumefaciens* as described in the previous section. Basta-resistant transformed plants on soil were selected using Glufosinate ammonium (AgrEvo). Plants with most reduction of *RGO* transcripts were chosen as silenced lines.

### Semi-quantitative RT-PCR and quantitative RT-PCR (qPCR)

Total RNA were isolated from rosette leaves, cauline leaves, stem and internode, and inflorescence apex tissues six-week old Arabidopsis WT plants grown under long-day conditions using Trizol reagent (Ambion, Thermo Fisher Scientific, Waltham, MA, USA). Total RNA from two-week old roots was isolated using RNAqueous® according to the manufacturer’s instructions (Ambion, Thermo Fisher Scientific, Waltham, MA, USA). Total RNA from young and mature siliques were isolated using a method described earlier (Murmu et al., 2010). RNA samples were treated with TURBO DNase™ (Ambion) prior to cDNA synthesis. Total cDNA from each RNA sample was synthesized from 2 μg of RNA template in a 20 μl reaction using Multiscribe reverse transcriptase (Applied Biosystems, Burlington, ON, Canada). The synthesized cDNA samples were diluted 1:5 with diethylpyrocarbonate (DEPC)-treated water (Ambion). Semi-quantitative RT-PCR was performed using 2 μl of diluted cDNA as template and gene-specific primers (Supplemental Table S3). Similarly, quantitative RT-PCR (qPCR) reactions were performed using 2 μl of diluted cDNA as the template and gene-specific primers in triplicate by Power SYBR Green Kit and in a StepOne Plus Real-Time PCR System according to manufacturer’s instructions (Applied Biosystems) as described earlier (Murmu et al., 2014). For absolute quantification of *RGO* transcripts in different tissues by qPCR, a standard curve method was used where the *RGO* standard curve was created using the *RGO*:pENTR-D-Topo plasmid, and *RGO* transcripts were calculated accordingly. Comparative *C*_T_ (ΔΔ*C*_T_) was used for relative quantification of transcripts of *GLK1* with GLK1-F2 plus GLK1-R2 primer set; and *RGO* with RGO-QF3 plus RGO-QR3 primer set and normalized using two endogenous control genes, *eIF4A-1* (*At3g13920*), and *GAPC* (*At3g04120*). The data represents three biological replicates with three technical replicates of each. All the qPCR data were analysed with *P* < 0.05 as statistically significant value using the StepOne 2.1 software (Applied Biosystems). A list of the genes and primers used in RT-PCR and qPCR is found in Supplemental Table S2.

### Green callus formation assay

Surface sterilized and stratified seeds were grown in square petri dish (VWR International, Ltd, Quebec, Canada) on half MS salt plus 0.7 % (w/v) Agar plates in the dark at room temperature for five days. After five days, root portion was excised out with a razor blade in a sterile laminar flow hood and immediately transferred onto half MS plus agar plate supplemented with 0.005 μM 2,4-D (2,4-Dichlorophenoxyacetic acid), a synthetic auxin, plus varying concentrations (0.005 to 0.5 μM) of BA (6-Benzylaminopurine) according to a procedure described earlier (Kiba et al., 2005). Plates were grown for an additional 30 days under long-day photoperiod conditions for generation of green calli. Green calli images were photographed with a Nikon D90 DSRL.

### Root growth assays

Sterilized seeds were placed on square petri plates (VWR International, Ltd, Quebec, Canada) containing minimal agar media (Haughn and Somerville, 1986) alone or supplemented either with 1% sucrose or 5 μm Kinetin (a synthetic CK). Plates were sealed with two layers of parafilm and grown vertically under a long-day growth cabinet for seven-day. At least 10 seeds from each genotype were placed on each plate with five to six replicates of the plates. The root growth assay was repeated three times with similar results. Roots were photographed with a Nikon D90 DSRL camera and root lengths were measured from the digital photographs with ImageJ software (https://imagej.nih.gov/ij/).

### Subcellular localization studies

For subcellular localization, leaf discs were prepared from three-week old soil grown Arabidopsis plants and stained for 1 hour with 10 μM DAPI (4’,6-Diamidino-2-Phenylindole, Dihydrochloride) in the dark at room temperature. After 1 hour the leaf discs were washed three times with sterile water and then mounted in Fluoromount-G ™ (Electron microscopy Sciences, Hatfield, PA, USA). Leaf discs were imaged with a Zeiss LSM800 Airyscan laser scanning confocal microscope (Carl Zeiss MicroImaging, Göttingen, Germany). For visualisation of DAPI and YFP, excitation lasers at 405 nm, 488 nm respectively were used and emission was monitored between 400 nm to 580 nm, 490 nm to 585 nm were acquired using the GaAsP detector and a Plan-Apochromat 63X/1.46 objective lens. For plasma membrane visualisation, leaf discs were stained with the lipophilic styryl fluorescent dye FM4-64FX (Molecular Probes, Eugene, OR, USA) at a concentration of 0.01 μg/mL in 1xPBS (Phosphate buffer saline) pH 7.4 and were incubated for 2 hours at room temperature in the dark. Subsequently, leaf discs were mounted in fresh dye and imaged with a Plan-Apochromat 63x/1.4 objective lens and the high resolution Airyscan detector using excitation laser line 488 nm and emission bands 490 nm to 580 nm, and 635 nm to 700 nm for YFP and plasma membrane, respectively. For endoplasmic reticulum (ER) localization of 35S:*GFP:HDEL*, an ER marker (Batoko et al., 2000) and 35S:*RGO:YFP* were infiltrated into four-week old *Nicotiana benthamiana* leaves via *Agrobacterium tumefaciens* according to a procedure previously described (Kosma et al., 2014). GFP and YFP fluorescence was monitored three days post infiltration. Essentially, leaf discs were mounted in Fluoromount-G ™ and YFP fluorescence was captured as described above. Similarly, GFP fluorescence was detected with excitation laser 488 nm and emission was monitored between 490 nm to 585 nm using the GaAsP detector and a Plan-Apochromat 63X/1.46 objective lens. For ER localization, similar plane and cell-types were imaged. All the confocal images were processed with ZEN 2.1 (Zeiss MicroImaging, Göttingen, Germany) and Adobe Photoshop CS6 (http://www.adobe.com/).

### Chlorophyll retention of dark-induced senescence with BA (6-Benzylaminopurine) treatment

Rosette leaves number 4 and 5 from four-week old plants of WT, 35S:*RGO*, and *rgo* plants grown in a long-day photoperiod conditions were used for this assay. Two set of six leaves from each genotypes were floated in a volume of 20 ml sterile water with 0.01N NaOH (−BA) or with 5 μM BA in 0.01N NaOH (+BA) in a petri dish, wrapped with aluminum foil and incubated in dark for seven days according to a procedure described earlier (Vercruyssen et al., 2015). After seven days of dark treatments (−BA, +BA), the leaves were imaged with a Nikon D90 digital SLR camera. The second set of leaves with similar treatments were used for chlorophyll (Chl: Chl a + Chl b) measurements. Two of 0.5 cm^*2*^ leaf disc were resuspended in 1ml DMF *(N,N-*Dimethylformamide) in the dark at 4° C overnight for total Chl extraction and Chl was measured with UV-visible spectrophotometer (Thermo Scientific) at 647 nm and 664.5 nm in 1 mm cuvettes. Total Chl was calculated per cm^2^ according to a procedure described earlier (Inskeep and Bloom, 1985). The experiment was repeated three times.

## ACKNOWLEDGEMENTS

We are grateful to Professor Tatsuo Kakimoto (Precursory Research for Embryonic Science and Technology, Japan Science and Technology Agency, Kawaguchi, Japan) for generously providing seeds of the Arabidopsis *ipt1 3 5 7* quadruple mutant and Professor Owen Rowland, Carleton University, for providing the GFP-HDEL construct.

## Notes

**Funding information:** This research was supported by the Genomics Research and Development Initiative (GRDI) of Agriculture and Agri-Food Canada to JS and RS

